# Scalable early detection of grapevine virus infection with airborne imaging spectroscopy

**DOI:** 10.1101/2022.10.04.510827

**Authors:** Fernando E. Romero Galvan, Ryan Pavlick, Graham Trolley, Somil Aggarwal, Daniel Sousa, Charles Starr, Elisabeth Forrestel, Stephanie Bolton, Maria del Mar Alsina, Nick Dokoozlian, Kaitlin M. Gold

## Abstract

Viral diseases, including Grapevine Leafroll-associated Virus Complex 3 (GLRaV-3), cause $3 billion in damages and losses to the United States wine and grape industry annually. GLRaV-3 has a well-studied, year-long latent period in which vines are infectious but do not yet display visible symptoms, making it an ideal model pathosystem to evaluate the scalability of symptomatic and asymptomatic imaging spectroscopy-based disease detection. Plant disease causes physiological and chemical changes to occur locally and systemically throughout a plant, which imaging spectroscopy can detect both directly and indirectly. Reliable and scalable disease detection during the latent period would greatly reduce management costs, as current detection methods are entirely ground-based, labor-intensive, and expensive. Here, we use data collected in September 2020 by the NASA Airborne Visible/Infrared Imaging Spectrometer Next Generation (AVIRIS-NG) to detect GLRaV-3 in Cabernet Sauvignon grapevines in Lodi, CA. During September 2020 and 2021, industry collaborators scouted 317 acres of Vitis vinifera winegrapes for visible disease symptoms, and collected a subset for confirmation molecular testing at a commercial facility. Grapevines identified as visibly diseased in 2021 were assumed to have been latently infected (asymptomatic) during the September 2020 AVIRIS-NG data collection. We combined random forest with synthetic minority oversampling technique (SMOTE) to train multiple spectral models able to distinguish between non-infected (NI) and GLRaV-3-infected grapevines. We observed clear spectral differences that allowed for differentiation between NI and GLRaV-3 infected vines both pre- and post-symptomatically at 1m through 5m resolution. Our two best performing models had 87% accuracy (0.73 Kappa) distinguishing between NI and asymptomatic (aSy), and 85% accuracy (0.71 Kappa) distinguishing between NI and (aSy + symptomatic [Sy]) respectively. We hypothesize these spectral differences are linked to changes in overall plant physiology induced by disease, as visible foliar symptoms were restricted to the lower canopy.

**Highlights:**

- Airborne imaging spectroscopy allows for scalable early-detection models of grapevine leafroll-associated virus complex 3 (GLRaV-3).
- Random Forest based models trained with scouting ground data and imaging spectroscopy are accurate up to 5 meter but perform best at 3 meter spatial resolution.
- GLRaV-3 detection via imaging spectroscopy will not replace existing field scouting strategies or molecular testing but supplement by allowing for more strategic resource deployment to improve the overall financial, environmental, and societal sustainability of winegrape production.

## 1. Introduction

Plant-microbe interactions can impact a variety of plant traits that can be remotely sensed, ranging from changes in tissue color to canopy architecture (1). Broadband methods relying primarily on visible (VIS) and nearinfrared (NIR) spectral indices, such as normalized difference vegetation index (NDVI), were proven capable of sensing late-stage plant disease in the 1980s (2; 3). However, the advent of more widely available, narrowband data streams spanning the VIS-shortwave infrared (SWIR) has revolutionized the study of plant disease sensing. Plant pathogens damage, impair, and/or alter foliar function, thus changing the chemical composition of foliage via the production of either systemic effectors or secondary metabolites, or by the physical presence of pathogen structures, such as hyphae and spores (4). These changes can be sensed with in situ and imaging spectroscopy, often referred to by plant pathologists as “hyperspectral imagery,” (5; 6; 7).

SWIR wavelengths have proved valuable for plant-pathogen interaction sensing due to their sensitivity to a range of foliar properties (8) including nutrient content (9; 10; 11; 12), water (13), photosynthetic capacity (14), physiology (15), phenolics and secondary metabolites (16; 17; 18), that are all impacted by early-stage disease. Recent work has established that airborne spectroscopic imagery can be used for pre-symptomatic disease detection in multiple pathosystems, including Pierce’s Disease of olive caused by Xylella fastidiosa (19; 20), Phytophthora spp. infection in Holm oak (21), and Oak wilt caused by the fungus Bretz (22; 23). These collective works showed that not only can airborne sensing provide reliable asymptomatic disease detection, in some instances it outperformed human scouting by detecting minimally symptomatic trees missed by the ground team (19).

Disease detection via imaging spectroscopy is often facilitated by machine learning, which helps to make sense of the underlying relationships within and amongst the hundreds of (highly correlated) spectral bands spectroscopic imagery offers (24; 25). Machine learning methods such as random forest and partial-least squares, have been used widely in airborne plant-microbe interaction sensing, including mycorrhizal association mapping (26), oak wilt detection (23), and Pierce’s Disease mapping (27; 28; 20), as well as in the broader, adjacent domains of foliar functional ecology (29; 30; 31).

While the above studies have shown imaging spectroscopy can be useful in understanding, detecting, and mapping plant-fungal and bacterial-fungal interactions, viral-plant interactions remain to be explored. Viral diseases, including that caused by Grapevine Leafroll Virus Complex 3 (GLRaV-3), cause three billion (USD) in damages and losses to the US wine and grape industry annually (32). GLRaV-3 is primarily vectored by mealybugs (Pseudococcidae sp.) but can also be propagated by other phloem feeding insects (33; 34; 35; 36). In addition to significantly reducing vine lifespan, GLRaV-3 infection causes the grapevine to misappropriate resources, which results in uneven cluster ripening, changes to grape berry chemistry, and reduced wine quality (32; 37). Existing strategies to detect GLRaV-3 in the field are based on visual scouting by trained experts (38). However, GLRaV-3 is particularly vexing to manage because only red grape varieties, (e.g. Cabernet Sauvignon, as opposed to Sauvignon Blanc) will ever display foliar symptoms for which humans to scout (39). Compounding this is the fact that GLRaV-3 has a long, approximately 12 month, latent phase during which the host is infectious but foliar symptoms are not yet apparent (32; 34; 40). This means that both latently infected red grape varieties and infected white variety grapevines serve as inoculum sources for nearby fields without grower recourse other than expensive molecular testing. Commercial, lab-based serological testing capable of identifying latent infections costs between $40-300 USD per vine depending on how many viruses are tested for and whether composite sampling is utilized. Even a small scale vineyard will have at least 1,000 vines, with larger vineyards containing up to 30,000, making both regular and asymptomatic testing impossibly costly to scale.

Consequently, plant pathologists and grape growers have begun to look for a detection approach that is both temporally and spatially scalable, accurate, and cost-effective. Remote sensing’s capacity for scalable, passive disease monitoring makes it of great interest to the plant pathology and broader agricultural science communities. Proof of concept work has established that contact and proximal imaging spectroscopy can detect GLRaV-3 infection at an early stage (41; 42; 43; 44). However, it has yet to be studied whether this capacity scales to aerial and airborne deployment scales. The goal of this work was to evaluate the scalability of airborne imaging spectroscopybased detection of symptomatic and asymptomatic grape viral disease with NASA’s Airborne Visible/Infrared Imaging Spectrometer Next Generation (AVIRIS-NG). In so doing, we sought to address the following questions: 1) How well does airborne imaging spectroscopy differentiate between non-infected and GLRaV-3 infected vines?, 2) Can detection be improved with various dimensionality reduction techniques? and 3) What is the optimal resolution for GLRaV-3 detection between 1-5m?

## 2. Data

Industry collaborators coordinated a team of highly trained field technicians to visually inspect (“scout”) 268 acres of red grape variety Aglianico (7ac), Cabernet Sauvignon (204ac), and Petite Sirah (57ac) for visible, foliar symptoms of GLRaV-3 according to industry best practices (38). Scouting and geotagging of visibly diseased vines were conducted in September, during harvest when symptoms are most apparent, in both 2020 and 2021. In total, 1427 and 2398 GLRaV-3-infected vines were identified in 2020 and 2021 respectively. In 2020, all grape clusters, green foliage, and canes were removed from the grapevines during mechanical harvest within one week of the final AVIRIS-NG flight. Cane tissue samples from 100 vines variably identified as diseased/non-diseased were sent to Agri-Analysis Laboratories (Davis, CA) for confirmation testing to validate scouting accuracy in 2020 and 10 samples in 2021. All samples sent for testing that had been identified as GLRaV-3-infected by the scouts returned as positive for infection. No samples that were identified as non-infected tested positive, giving us high confidence in the scouting teams’ accuracy. In between the 2020 and 2021 growing seasons, our industry collaborators removed diseased vines from the vineyards after harvest and prior to the following season’s bud break to prevent them from serving as an inoculum reservoir for uninfected grapevines. This means that vines identified as diseased in 2020 were not present in the field in 2021.

Imagery from the NASA Jet Propulsion Laboratory AVIRIS-NG was acquired in September 2020. This campaign yielded data over 37,317 acres of California vineyards at the peak of the growing season. Each acquisition collected submeter spatial pixels in 425 evenly-spaced bands sampling the 380nm to 2510nm spectral range at 5nm intervals (45; 46). Acquisitions were collected between 1:00 and 3:00 PM Pacific (local) time. A subset of this total imagery collected over 204 acres of Cabernet Sauvignon grapevines in Lodi, California on September 18th were used for this study (Figure 5). Specifically, flightlines used for this work are: ang20200918t210249, ang20200918t205737, ang20200918t212656, ang20200918t213801, ang20200918t213229. All AVIRIS-NG imagery used in this study is reflectance data and is publicly available and can be downloaded from the AVIRIS-NG data portal https://aviris.jpl.nasa.gov/dataportal/.

Generic spectra for soil, vegetation, and shadow were used as endmembers for spectral-unmixing. The soil and vegetation spectra were pulled from the United States Geological Survey (USGS) Spectroscopy Lab’s spectral library (47). These spectral measurements were selected because they match the spectral range of the AVIRIS-NG imagery in terms of spectral resolution. Additionally, the Lodi team provided ground-spectral measurements of grasspastures, soil, pavement, and vines for calibration and validation purposes (Figure Supp-3).

## 3. Methodology

The entirety of the pipeline outlined in Figure 1 was written in Python 3.9. All scripts are available on the GoldLab Github repository (https://github.coecis.cornell.edu/GoldLab-GrapeSPEC). Anaconda was used for python package management. Likewise, anaconda virtual environments have been uploaded to GoldLab-GitHub.

**Figure 1:**
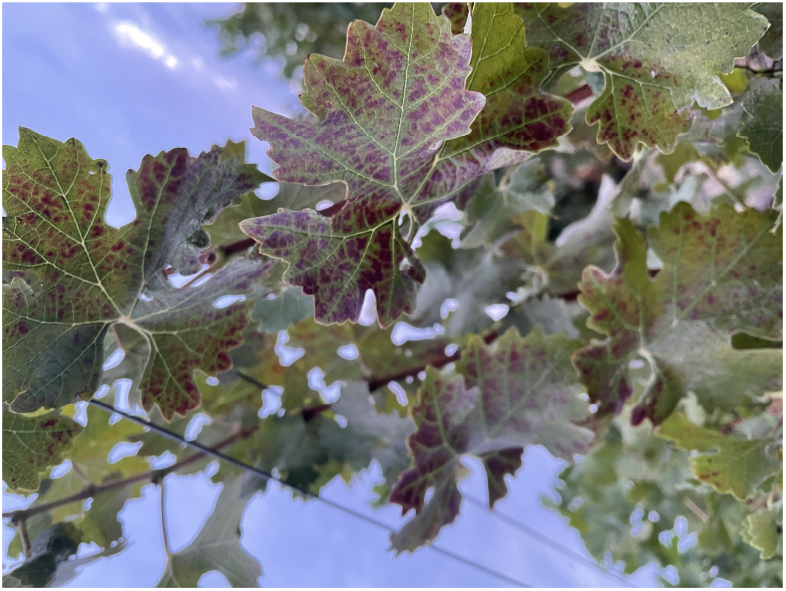
GLRaV-3 infected grapevine of Cabernet Sauvignon variety.

### 3.1. Preprocessing AVIRIS-NG Imagery

Both bidirectional reflectance distribution function (BRDF) and topographic correction (48) were applied to all reflectance files of the AVIRIS-NG imagery. BRDF and topographic correction code were pulled from the Hy-Tools package (49), an open-source spectroscopy processing python library. All flight lines were corrected in full. Some of the AVIRIS-NG imagery required further spatial georeferencing to be better aligned with the disease incidence coordinates collected by the scouting team. National Agriculture Imagery Program (NAIP; USDA) imagery collected within a week of our AVIRIS-NG imagery was used as a reference to improve georeferencing and co-registration. Each NAIP image was clipped according to the area of overlap with the AVIRIS-NG. Specifically, the NAIP red band was extracted and compared to the AVIRIS-NG band at 600nm. Ground control points (GCPs) were generated by passing the described target and reference imagery to the open-source python library Automated and Robust Open-Source Image Coregistration Software (AROSICS) (50); To be even more specific, AROSICS was parameterized according to the code-base author’s default parameters, after processing and a quick visual inspection, we found the image was successfully co-registered. The resulting GCPs were then used to co-register the AVIRIS-NG imagery onto the NAIP imagery. Lastly, following standard practice, noisy bands (due to water absorption and other atmospheric effects) present in the data were excluded following a visual analysis; these are the wavelengths between 380 - 400nm, 1310 - 1470nm, 1750 - 2000nm, and 2400 - 2600nm, these removed wavelengths are illustrated as gaps in Figure 3 and Supplementary Figures 7 and 8.

Vineyard boundaries shared by our industry collaborators were used to mask the AVIRIS-NG imagery. The open-source Python library Rasterio’s mask package was used for clipping the imagery (51). The aforementioned open-source HyTools library included a command line interface (CLI) python spatial resampling script that was used to resample the clipped co-registered imagery from the native one-meter (1m) to three-meter (3m) and five-meter (5m), the spatial resampling script applies a nearest neighbor algorithm with no blurring effects to resample the image.

The spectral mixture residuals (MR) was computed using the open-source code of (52). Endmembers for shadow, soil, and generic vegetation were used to spectrally unmix the AVIRIS imagery (Figure 6). The SMR simultaneously estimates two related parameters for each pixel in the image: 1) fractional area of each input endmember, and 2) mixture residual spectra quantifying wavelength-explicit misfit. Soil masks for each vineyard were generated by calculating a percentage of vegetation and soil endmember weights calculated by the spectral MR script. Specifically, the formula used was Vw/(Vw + Sw); where Vw is the vegetation endmember fractal weight and Sw is the soil endmember fractal weight. The resulting raster was then used to create a binary mask, all pixels with a value ≥ 0.50 were assigned a one, and all remaining pixels were assigned a zero. The percentage of 0.5 was found to be the optimal percentage that both preserved disease incidence and removed mostly soil pixels. This binary mask was then applied to all clipped spectroscopic imagery retaining only pixels sampling a minimal area which was not covered with grapevine canopy.

### 3.2. Model Training Pipeline

#### 3.2.1. Dataset Labels

In total, our dataset contains 3 labels: non-infected (NI), symptomatic (Sy), and asymptomatic (aSy). GLRaV-3-infected grapevines of red grape variety can stay at the asymptomatic stage for up to a year if no visible symptoms show before a grapevine’s winter dormant period (34; 40). Therefore, given our understanding of disease biology, we label vines identified as visibly infected in 2021 as asymptomatic (aSy), as there is a high likelihood they were latently infected at the time of the 2020 AVIRIS-NG flight. Vines identified as visibly diseased by the scouts in 2020 are labeled as symptomatic (Sy). However, we caveat that we do not have molecular testing to prove that these vines were truly asymptomatically-infected at the time of data collection, as it would have been unfeasible given our scope of study and sample testing expense ($40-50 per vine). We emphasize that our assumption is well-supported by current understanding of disease biology (53; 54; 32) and the fact that all green foliage was destroyed and removed from the vineyard during mechanical harvest soon after the flight took place. This means it is unlikely that vines experienced an opportunity to become infected between the time of the AVIRIS-NG flight and bud break the following season, since green tissue is required for the insect-vector to feed and transmit disease.

To generate our “non-infected” vine locations, clipped, co-registered, and soil-masked imagery was vectorized by extracting the centroid for all pixels within the image. GLRaV-3 infections are known to cluster spatially (55; 56; 57), so centroids within 5m of the known diseased coordinates were excluded to avoid accidentally labeling vines as “non-infected” that may not truly be non-diseased. In total 621,000, 70,000, and 27,000 non-infected vine pixels were found using the outline above for 1m, 3m, and 5m resampling imagery respectively. The pixel count for aSy were 2258, 2458, and 2192 for 1m, 3m, and 5m. The pixel count for Sy were 1027, 1139, 1031 for 1m, 3m, and 5m. The pixel count for the class where aSy and Sy were treated as one class was 3285, 3597, 3223 for 1m, 3m, and 5m.

#### 3.2.2. Random Noise and Dimensionality Reduction

We evaluated multiple approaches to reduce noise in our spectroscopic imagery. The first strategy involved using the SMR output as the features for the RF to train on. The second strategy was using the Savitzky-Golay filter (SG) from the signal module in the Scipy python package (58). This SG smoothing window was limited to only five wavelengths and polynomial order of three. Thirdly, SKLearn’s PCA library was used to perform a principal components analysis (PCA) on the spectral feature space to reduce the dimensionality of the feature space down from 425 to 10 principal components (PCs). Lastly, a combination of all techniques above were used as the feature set for the later training of the RF model. All noise-reduction and dimensionality reduction techniques excluded noisy wavelengths specified in section 3.1.

#### 3.2.3. Spatial Sampling

Coordinates for symptomatic, asymptomatic, and non-infected vines were loaded onto one Geopandas data frame. All coordinates were stored and reprojected to the AVIRIS-NG acquisition’s coordinate system WGS 84 / UTM zone 10N; EPSG: 32610. Each coordinate was then used to spatially sample the AVIRIS-NG imagery, resulting in a data frame that held the coordinate, label, unique field name, and relevant spectral data.

#### 3.2.4. Balancing Classes

Non-infected vines far outnumber diseased vines. In our data, non-infected:diseased ratios were approximately 3109:18, 700:33, 175:9 for 1m, 3m, and 5m respectively, which presented a challenge in choosing a balancing strategy. Specifically, the challenge was to select a representative population of non-infected vines to train on given the diversity within each wavelength in our spectroscopic imagery. Therefore, two main strategies were followed and compared: undersampling and oversampling.

To undersample, a PCA was performed on the spectra of all non-infected vines, in total, three PCs were found to explain 95% of the variance of the non-infected-vine spectra. The resulting PC space was then clustered using SKLearn’s K-Means clustering library, three K-Means were found to be optimal to cluster the data using the ‘elbow method’, where distortion is plotted against the number of clusters to establish a cut-off, in our case, three-clusters were found to be optimal. Next, an equal number of points were randomly selected from the resulting three-dimensional cluster feature space, in total 3200 NI spectral rows were subsampled from 1m, 3500 for 3m, and 3200 for 5m.

To oversample, Imbalanced Learn’s (59) Synthetic Minority Oversampling Technique (SMOTE) was used as our oversampling strategy. However, oversampling the disease incidence count (aSy and Sy) point count to the NI vine count is by no means an appropriate approach given that the disease incidence count sits at about 3,000 and NI at about 600,000. When we attempted to increase the disease incidence count to the NI count data-points close to duplicates were identified. Therefore a mixture of undersampling and oversampling was followed. First, the diseased incidence points were oversampled using SMOTE to increase their count by 10%, 25%, 50%, 75%, 100%, 200%, and 300%. We found an increase of 50% oversampling of the minority class best

### 3.3. Training Random Forest Models

Scikit Learn’s RF python package was used for RF model training (60). The now balanced datasets were used to train 10 RF models, and performance was averaged across the models for the validation set. The training/validation data was split 70/30 and 10 k-folds were done on each training. All steps were repeated for all transformations done to the spectra and across the 3m and 5m spatially resampled data.

## 4. Results and Discussion

### 4.1. Best Performing Model

All models produced in this study followed a 70/30 training and validation scheme. All accuracy and respective kappa scores listed in Table 1 are derived from the 30% validation data withheld during training. Lastly, each metric presented is the result of the average of 10 models’ performance. Overall, our two best performing models were for discrimination between 1) non-infected and asymptomatic infected vines (NI vs aSy) and 2) non-infected and a combined dataset of asymptomatic and symptomatically infected vines (NI vs [aSy+Sy]) resampled to 3m resolution with Savitzky-Golay and principal-components used as training data with a balancing strategy that first undersampled the majority class and oversampled via SMOTE (Table 1). These models respectively had 87% accuracy (0.73 Kappa) and 85% accuracy (Kappa = 0.71) in discriminating between non-infected and diseased (both visually and non-visually) vines. I Spectral transformations of various types were considered and tested, in large part, to reduce noise, dimensionality, and correlation between wavelengths. Savitzky-Golay (SG) is commonly used filtering methods used as a noise-reduction technique in signal processing and has a history of reducing random-noise from spectroscopic data (61). The logic behind using Savitzky-Golay (SG) was to reduce the noisiness of the continuous spectral signal by smoothing the spectro-scopic signal, SG by applying an SG filter with a small window and low order polynomial improving our RF model when compared to data with no smoothing. Using the spectral residuals as the feature set for training the random forest model improved performance than using unaltered reflectance values. Particularly, we found that the MR dataset improved differentiation between NI, Asy., and Sy vines when compared to reflectance or reflectance with smoothing via SG filter. However, we found that training a model with PCs generated via SG filtered reflectance outperformed the models trained with MR.

**Table 1:** Columns depict the transformation(s) applied to the spectral features. Partial Least Squares Discriminant Analysis (PLS-DA), Random Forest (RF), Savitzky-Golay (SG), Principal Component Analysis (PCA), Spectral Unmixed Residuals (SMR), Synthetic Minority Oversampling Technique (SMOTE). Mean accuracy of true label (A) and Kappa score (K).

| Res | Class | PLS-DA |  | RF |  | RF SG |  | RF SG PCA |  | RF MR |  | RF MR PCA |  | RF SG PCA SMOTE |  |
| --- | --- | --- | --- | --- | --- | --- | --- | --- | --- | --- | --- | --- | --- | --- | --- |
|  |  | A | K | A | K | A | K | A | K | A | K | A | K | A | K |
| **1m** | NI vs Sy+aSy | 74% | 0.46 | 70% | 0.40 | 72% | 0.44 | 75% | 0.51 | 72% | 0.45 | 73% | 0.45 | 78% | 0.56 |
|  | NI vs Sy | 68% | 0.34 | 63% | 0.26 | 65% | 0.29 | 65% | 0.30 | 67% | 0.38 | 63% | 0.24 | 68% | 0.36 |
|  | NI vs Sy vs aSy | 59% | 0.36 | 53% | 0.29 | 53% | 0.30 | 56% | 0.34 | 56% | 0.35 | 54% | 0.31 | 61% | 0.41 |
|  | NI vs aSy | 79% | 0.57 | 75% | 0.50 | 78% | 0.55 | 81% | 0.62 | 76% | 0.50 | 79% | 0.58 | 83% | 0.66 |
| **3m** | NI vs Sy+aSy | 75% | 0.46 | 68% | 0.38 | 75% | 0.48 | 73% | 0.46 | 76% | 0.48 | 75% | 0.48 | **85%** | **0.71** |
|  | NI vs Sy | 73% | 0.47 | 62% | 0.24 | 72% | 0.41 | 70% | 0.41 | 74% | 0.47 | 74% | 0.47 | 81% | 0.62 |
|  | NI vs Sy vs aSy | 62% | 0.43 | 48% | 0.23 | 57% | 0.34 | 57% | 0.35 | 61% | 0.49 | 60% | 0.39 | 67% | 0.51 |
|  | NI vs aSy | 75% | 0.49 | 70% | 0.41 | 80% | 0.58 | 83% | 0.66 | 77% | 0.51 | 76% | 0.49 | **87%** | **0.73** |
| **5m** | NI vs Sy+aSy | 76% | 0.51 | 70% | 0.41 | 73% | 0.46 | 77% | 0.56 | 77% | 0.52 | 75% | 0.46 | 80% | 0.59 |
|  | NI vs Sy | 74% | 0.47 | 60% | 0.21 | 69% | 0.38 | 71% | 0.41 | 75% | 0.47 | 72% | 0.40 | 75% | 0.51 |
|  | NI vs Sy vs aSy | 63% | 0.43 | 50% | 0.26 | 59% | 0.38 | 61% | 0.42 | 64% | 0.45 | 61% | 0.39 | 66% | 0.48 |
|  | NI vs aSy | 79% | 0.57 | 74% | 0.48 | 75% | 0.50 | 81% | 0.67 | 77% | 0.54 | 77% | 0.54 | 83% | 0.67 |

### 4.2. Scaling

Airborne spectroscopic imagery pixel size is dependent on the height the aircraft is flown at. Therefore, the exact AVIRIS-NG pixel size achieved can be unpredictable, though will commonly be between 1-5m. We sought to determine whether GLRaV-3 detection accuracy varies within this range. Overall, the accuracy of our models did not vary significantly across spatial scales ranging from native resolution (1m) to resampled 3m and 5m (Table 1). We observed that our random forest model performed best at 3m spatial resolution, at 5m and 1m, there is a decrease in accuracy. We suspect that the drop in accuracy between 3m to 5m is due to dilution of the spectral signal that underlies our ability to discriminate between groups. We anticipated that detection would be most accurate at native, 1m resolution, however we do not find that to be the case. We suspect that this is because each individual diseased vine, despite the geotag being accurate to the submeter, is sometimes at the edge of multiple-pixels, which may conflate the signal. Additionally, at one meter resolution there is less dilution of things that cause noise in the data, such as soil patches, grass, and other none-vine reflectance. We found 3m to be the optimal balance point between minimizing noise and not over-diluting the dataset.

### 4.3. Asymptomatic detection and class differentiation

In all the cases we evaluated, the diseased label, regardless of symptomatic or asymptomatic, was differentiable from the non-infected vine spectra (Figure 2). This finding that all three classes differ spectrally supports our assumption that our asymptomatic label vines were indeed experiencing latent infection at the time of data acquisition. When classifying between all three classes, we see the majority of model confusion seems to be between symptomatic and asymptomatic vines, indicating that these groups share enough spectral commonality between them for a model to disproportionately confuse them. We believe this spectral commonality is due to GLRaV-3-infection, which is known to impact vine biology prior to symptom appearance (42; 43; 44).

**Figure 2:**
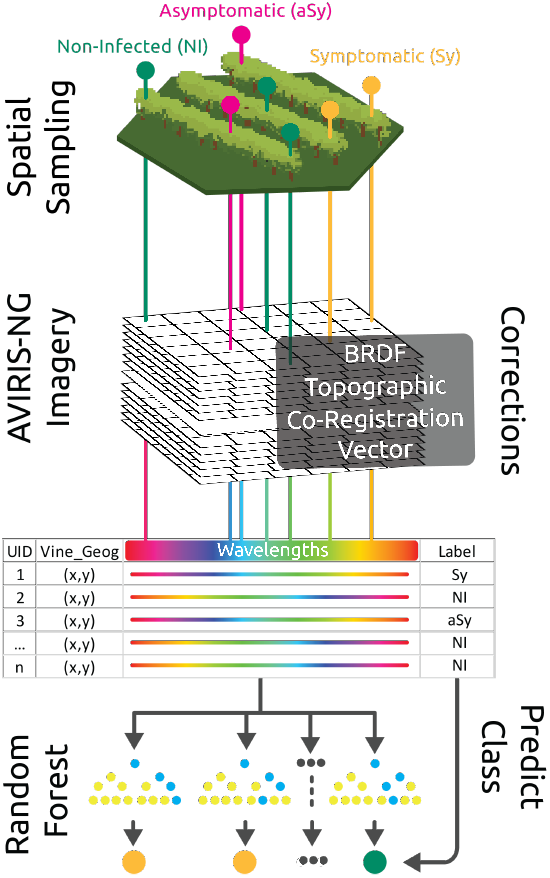
Illustrative summary of the pipeline.

Additionally, we observe non-infected and asymptomatic vines have different average reflectance from each other across the spectrum (Figure 3). A vegetation mask was used to generate a set of points for non-infected vines outside a 5m buffer of infected-vine locations. This method assumes that there is no GLRaV-3 at the asymptomatic stage at these locations, that each point is indeed a vine and not a patch of grass, or that the confirmed in-fected vine does not go beyond the buffer space. Increasing the accuracy of healthy vine location and more information regarding the status of each vine would undoubtedly yield a better performing model but would require a tremendous amount of resources to properly measure.

**Figure 3:**
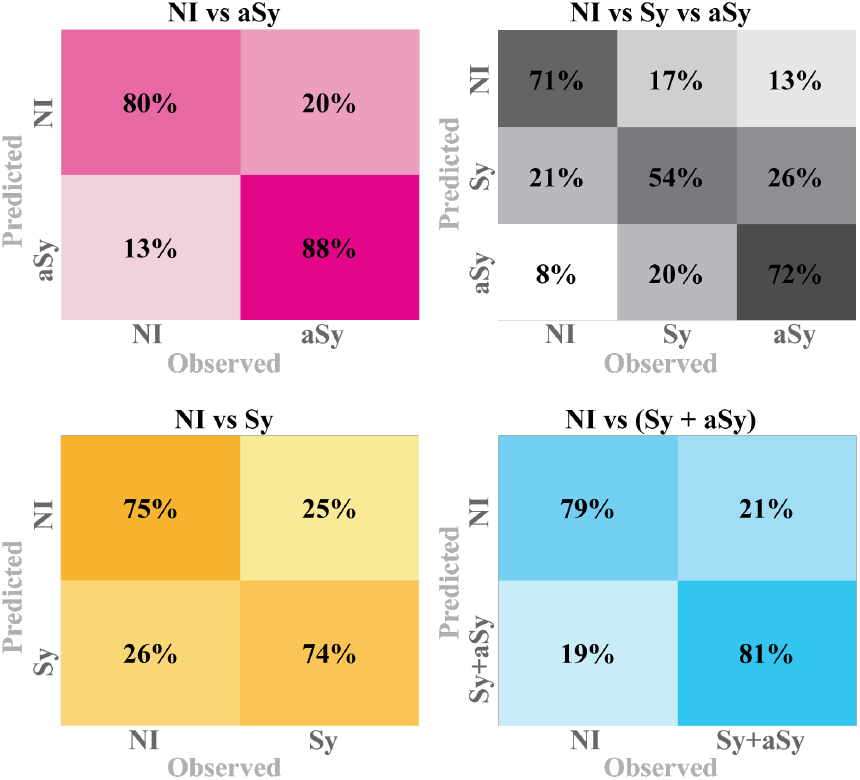
Confusion matrices on validation datasets for Random Forest 3m SG-PCA-SMOTE.

We used our 3m SG-PCA-SMOTE model to classify AVIRIS-NG imagery with the appropriate corrections and transformations to the data (Figure 4). In all cases, we allow the RF model to classify any pixel with a 50% probability as infected. We find that while overall, accurate, the model over-classifies areas as diseased (asymptomatic or symptomatic) than the true observed diseased extent. We observe that these misclassification swathes tend to cluster near the edges of the vineyard boundary. We suspect that the misclassifications here are due to confounding instances of biotic (disease) and abiotic stress. Vines at the ends of the row are known to experience more stress than buffered, inner vines. This includes more exposure to the elements and uneven management maintenance (e.g., irrigation being lesser near the row end). Untangling biotic and abiotic stressors is a major challenge to accurate plant-pathogen interaction mapping with imaging spectroscopy. This is compounded by the fact that viral diseased vines are generally more susceptible to the negative physiological impacts of abiotic stressors, since they have a less healthy foundation than an uninfected vine. Additionally, fields are not sterile environments, and it is likely that an infected vine may have a combination of stressors affecting it, such as pest damage. We suspect it likely that these border regions that are experiencing disproportionately more stress have more similar than not spectral profiles at this scale of study to infected vines. Alternatively, the confusion may be due to an overabundance of water. Proximity to water, such as bounding a river as our example vine-yard in Figure 4 does, can increase the likelihood of unhealthy root-systems and fungal infections in crops, especially for grapevines with deep, reaching roots.

**Figure 4:**
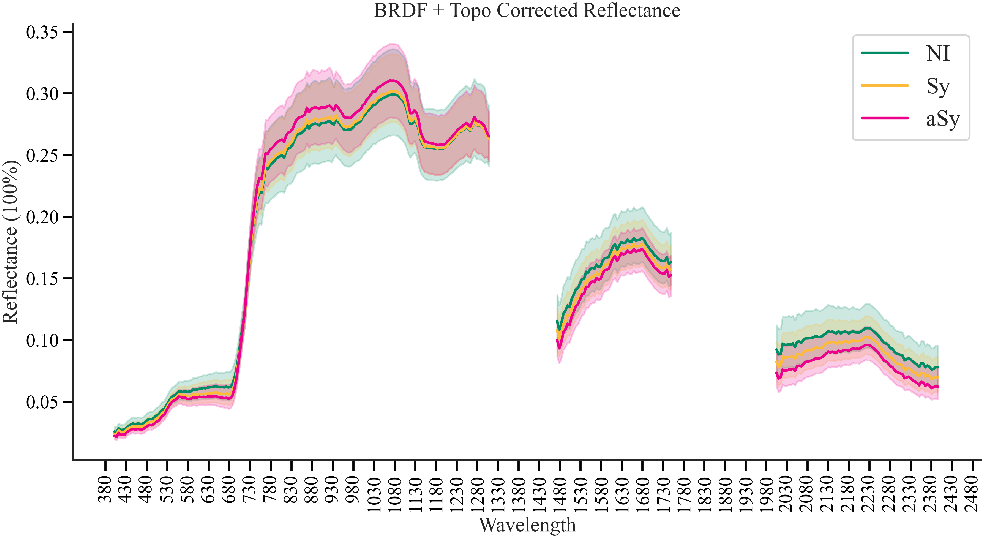
Reflectance values by vine status; Symptomatic (Sy), aSymptomatic (aSy), Non-Infected (NI).

**Figure 5:**
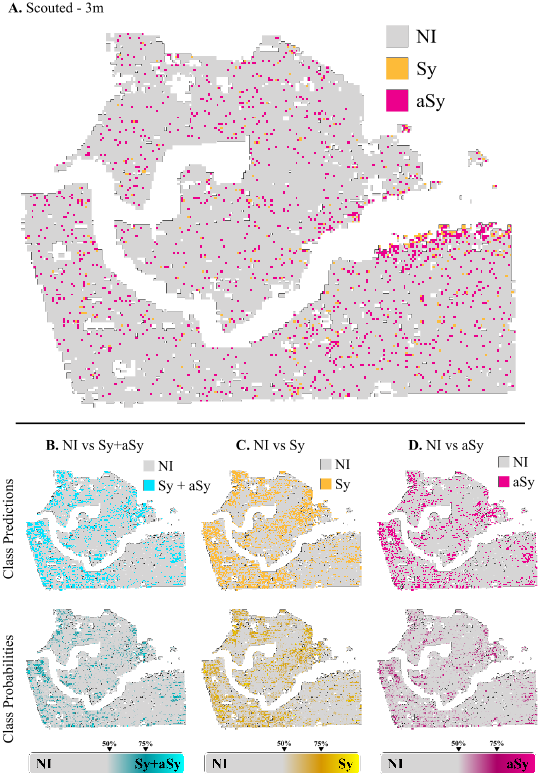
Classified AVIRIS-NG spectroscopic image over vineyard in California, Lodi using the RF model SG-PCA-SMOTE at 3m. (NI) - None-infected, (Sy) - Symptomatic vines, (aSy) - Asymptomatic vines. A. Scouted locations of GLRaV-3 at 3m resolution. B. Predictions of model trained to differentiate between NI and [Sy+aSy] and respective probability heat-map. C. Predictions of model trained to differentiate between NI and Sy and respective probability heat-map. Predictions of model trained to differentiate between NI and aSy and respective probability heat-map.

Indeed, one of the biggest challenges in disease detection via imaging spectroscopy is untangling the signal of abiotic and biotic stressors that may be affecting a vine simultaneously. Typical vineyard abiotic stressors, including those related to water-stress or nutrient deficiency, are closely related to thermal data (27; 62). Abiotic and biotic stresses, even those with a common visual manifestation (e.g. wilt) can be differentiated with airborne sensing because their underlying biological origins are different (19). Multi-modal sensing compliments imaging spectroscopy-based disease detection by adding biologically relevant data that can be useful for untangling biotic and abiotic stress. In other pathosystems, including solar induced fluorescence (SIF) has proven useful given the close relationship between SIF and photosynthetic activity (63). However, calculating SIF requires a narrower band instrument (> *3nm*) than that offered by AVIRIS-NG (5nm). The addition of thermal and SIF products in combination with spectral unmixing approaches, such as SMR or Multiple Endmember Spectral Mixture Analysis (MESMA) could help provide a clearer picture of the biological origins, biotic or abiotic, of crop stress. Despite these challenges, we still find that spectroscopic imagery mapping is a rapid and accurate GLRaV-3 detection tool.

## 5. Conclusions

Interest in using non-destructive imaging spectroscopy to detect plantmicrobe interactions has grown exponentially in recent years, however most explorations have focused on bacterial and fungal disease detection in tree crops (20; 26; 23). Our work expands our current understanding of plant disease sensing by reporting for the first time the capacity for airborne spectroscopic imagery to detect both symptomatic and asymptomatic plant-viral interactions in a non-tree crop at multiple resolutions. Our findings suggest that viral infection, regardless of visible symptom appearance, imparts a consistent, systemic change to foliar reflectance that can be detected with airborne imaging spectroscopy, and that this ability is improved by de-noising and dimensionality reduction techniques. A scalable, non-destructive, and low-cost solution for asymptomatic viral infection detection is a game changing prospect for the grape industry and agriculture at large. Destructive molecular or serological testing remains the most accurate method to detect viral infection at the asymptomatic stage, however this approach is impossible to scale due to expense. The methodology we offer here serves as the initial steps for accurate and scalable early detection of grapevine viral infection that could be used to deploy ground mitigation efforts, such as scouting, molecular testing, and vine removal, more strategically.

Geography plays an important role in generalizability of all remote sensing applications. The models developed here only consider central California where the soil type, grape variety, regional vineyard management practices, and climate likely impact model performance. Future work should investigate an expanded geographic range, such as different latitudes of California, and eventually, outside of California such as the northeastern great-lakes region, or other wine-producing countries. Additionally, crop varieties are known to interact with pathogens differently (5). In grapevine, for example, white grape varieties (such as Chardonnay) display extremely subtle to no foliar symptoms when infected with GLRaV-3, and some hybrids are known to have higher viral load tolerance levels that impacts biological response (32). In grape production, varieties are managed differently according to their individual requirements as well as the product (e.g. wine, juice, table, or raisin) a grower is aiming to deliver. These differences all influence spectral signals. Scientists aiming to use spectroscopic imagery for disease detection not only in grapevine, but any crop system, must take into consideration generalizability along varieties, geography, and management practices and quantify how each change affects disease-detection.

Using airborne imaging spectroscopy and machine learning, we have developed models that are effective at identifying spectroscopic signal of GLRaV-3 at various spatial resolutions. Our work does not aim to replace existing field scouting strategies or molecular testing. Rather, our work paves a path forward for more strategically deploying these resources to improve the overall financial, environmental, and societal sustainability of winegrape production. The next steps for this work are to assess scalability to space-borne resolution for use with NASA’s forthcoming Surface Biology and Geology satellite. Our long term goal for this project is to assess the space-borne scalability of our findings, and to develop it such that it is readily available to the grape and broader agricultural community in such a form that little programming, remote sensing, or GIS expertise would be required to use them. For this effort, a cloud-based solution with intuitive front-end design or edge-computing deployment is recommended.

## 6. Declaration of Competing Interests

The authors declare to have no conflicts of interest/competing interests.

## 7. Acknowledgements

This work received funding from NASA FINESST grant number 80NSSC21K1605, the NASA Jet Propulsion Laboratory Strategic University Research Partnership Fund, and the NASA Biodiversity and Ecological Forecasting program office. Romero Galvan received funding from the NSF NRT Digital Plant Sciences grant awarded to Cornell University (Grant number 1922551). Sousa gratefully acknowledges funding from the USDA NIFA Sustainable Agroecosystems program (Grant number 2022-67019-36397), the NASA Land-Cover/Land Use Change program (Grant number 21-LCLUC21_2-0025), the NASA Remote Sensing of Water Quality program (Grant number 80NSSC22K0907), and the NSF Signals in the Soil program (Award number 2226649). We acknowledge with gratitude the role of Michael Eastwood and the many members of the NASA JPL AVIRIS-NG team who, along with Woody Turner, are responsible for initiating the successful air campaign that led to this publication. Importantly, we would like to thank our industry collaborators for their invaluable efforts and collaboration in supporting this project. In particular, we thank the talented and hardworking scouting teams who spent countless hours in vineyards identifying and geotagging diseased vines to provide us with high quality validation data.

## 8. Supplementary material

**Supplementary Figure 6:**
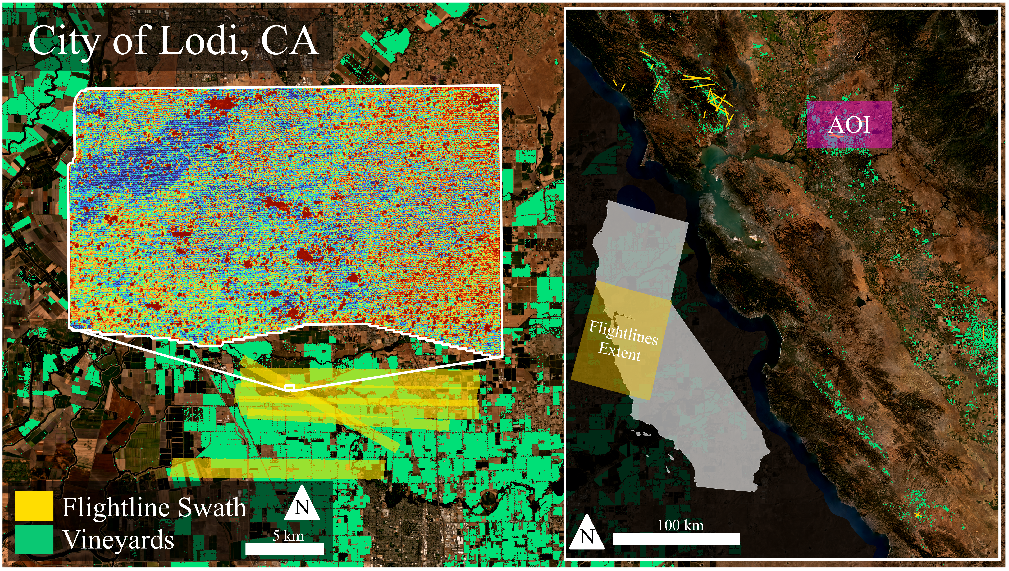
Region of interest, vineyards overlapping AVIRIS-NG spectroscopic imagery collected over the city of Lodi, CA.

**Supplementary Figure 7:**
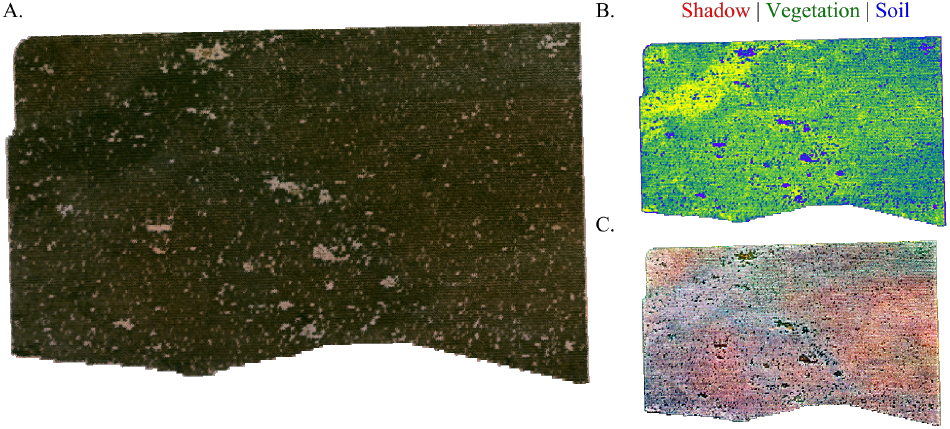
A. corrected AVIRIS-NG Reflectance image at one-meter resolution, B. Spectrally unmixed weights as RGB, C. Spectrally unmixed residuals.

**Supplementary Figure 8:**
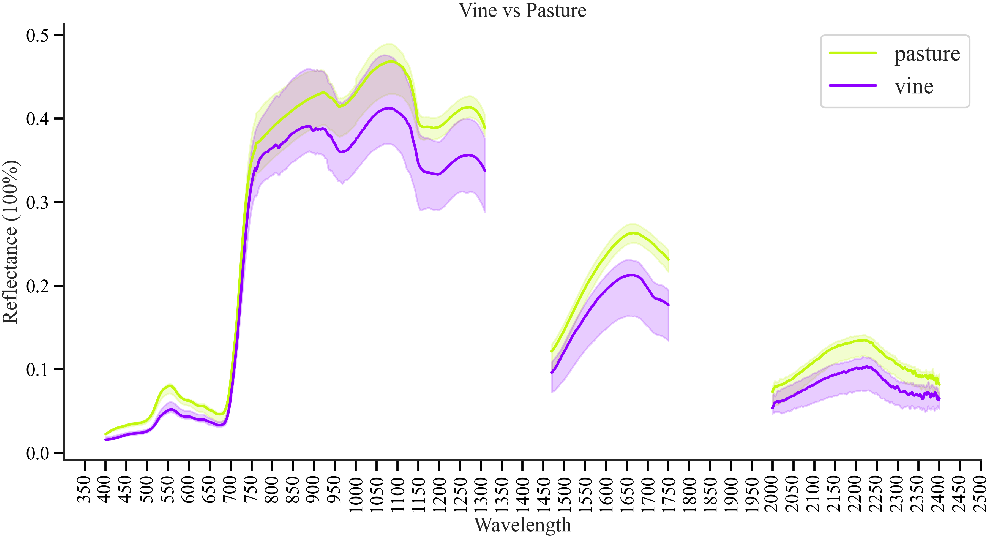
Reflectance of vines and pasture in Lodi vineyard taken by hand-held spectrometer.

**Supplementary Figure 9:**
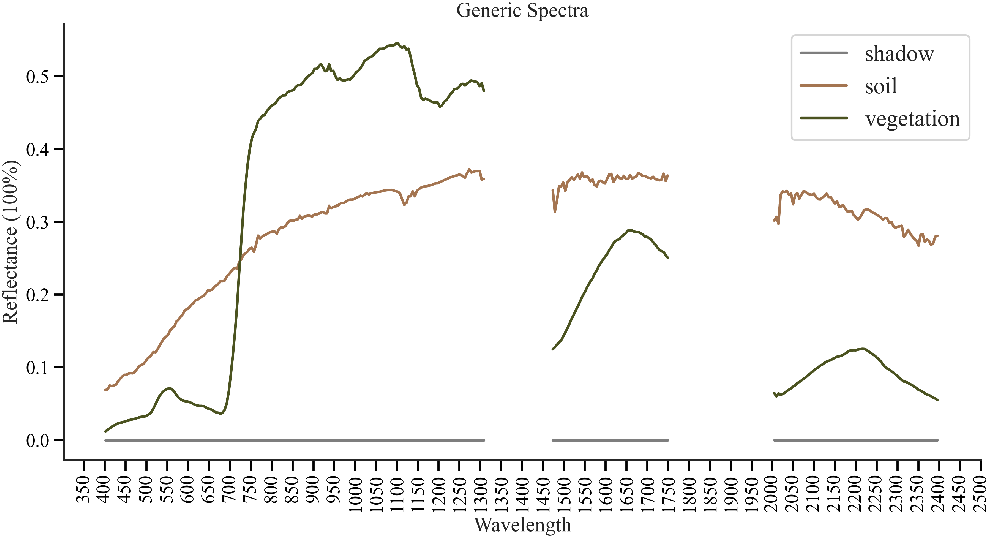
Reflectance of generic spectra for vegetation, soil, and shadow displayed as photometric zero.

## Notes

### Competing Interest Statement

The authors have declared no competing interest.

